# Three-dimensional chromatin mapping of sensory neurons reveals that cohesin-dependent genomic domains are required for axonal regeneration

**DOI:** 10.1101/2024.06.09.597974

**Authors:** Ilaria Palmisano, Tong Liu, Wei Gao, Luming Zhou, Matthias Merkenschlager, Franziska Müller, Jessica Chadwick, Rebecca Toscano Rivolta, Guiping Kong, James WD King, Ediem Al-jibury, Yuyang Yan, Alessandro Carlino, Bryce Collison, Eleonora De Vitis, Sree Gongala, Francesco De Virgiliis, Zheng Wang, Simone Di Giovanni

## Abstract

The in vivo three-dimensional genomic architecture of adult mature neurons at homeostasis and after medically relevant perturbations such as axonal injury remains elusive. Here we address this knowledge gap by mapping the three-dimensional chromatin architecture and gene expression programme at homeostasis and after sciatic nerve injury in wild-type and cohesin-deficient mouse sensory dorsal root ganglia neurons via combinatorial Hi-C and RNA-seq. We find that cohesin is required for the full induction of the regenerative transcriptional program, by organising 3D genomic domains required for the activation of regenerative genes. Importantly, loss of cohesin results in disruption of chromatin architecture at regenerative genes and severely impaired nerve regeneration. Together, these data provide an original three-dimensional chromatin map of adult sensory neurons in vivo and demonstrate a role for cohesin-dependent chromatin interactions in neuronal regeneration.

## INTRODUCTION

Hi-C studies have revealed that the genome is folded into self-interacting regions known as topologically associating domains (TADs) or contact domains, which regulate gene transcription by spatially restricting and facilitating enhancer-promoter interactions (1–4). The three dimensional (3D) chromatin organization allows enhancers to loop over long genomic distances to engage with target gene promoters, affecting gene transcription by providing binding sites for transcription factors (TFs) and chromatin remodelers, and by recruiting RNA polymerase II machinery and RNA polymerase II regulators (5, 6). Such 3D organization arises from the coordinated activity of cohesin and CCCTC-binding factor (CTCF) (7–10). According to the loop extrusion model, the ring-like cohesin complex translocates on the DNA in an ATP-dependent way progressively extruding bidirectional chromatin loops until it is halted by CTCF bound in a convergent orientation (11, 12).

Despite the critical role of 3D genome looping in gene regulation, the in vivo map of chromatin architecture in mature adult neurons at homeostasis and after injury remains elusive. Similarly, whether chromatin organization and enhancer-promoter interactions are modified by loss of neuronal homeostasis and whether these interactions play a role in axonal regeneration after injury remain undetermined. The axonal regenerative capacity in the peripheral nervous system is underpinned by the coordinated changes in the expression of hundreds of genes involved in multiple interconnected biological processes (13–15). Studies in the dorsal root ganglia (DRG), which contain sensory neurons, have shown that histone acetylation, DNA methylation and hydroxymethylation contribute to the regenerative transcriptional program after nerve injury by affecting chromatin accessibility at promoters and enhancers of regenerative genes (15–24). However, they did not show an absolute requirement of these epigenetic mechanisms for nerve regeneration, and they did not account for the role of 3D chromatin folding in the regulation of gene expression after injury. Furthermore, when systemic studies were performed, they were conducted from bulk DRG tissue and did not discriminate between neurons versus stromal and glial cells (15, 25, 26).

Here we provide a chromatin architecture map of *in vivo* mature sensory neurons and determine the role of neuronal 3D chromatin organisation in gene expression regulation at homeostasis and after a nerve injury. To this end, we performed Hi-C and RNA-seq in purified DRG neurons following sciatic nerve crush (SNC) from wild-type (WT) and *Rad21* (cohesin structural subunit) conditional knock-out (KO) mice. We show that 3D chromatin integrity of regenerative genes is required for nerve regeneration. Cohesin is needed for the organization of 3D genomic domains indispensable for the full activation of regenerative genes. Importantly, loss of cohesin prevents nerve regeneration *in vivo*. These data provide an *in vivo* neuronal 3D chromatin and gene expression map at homeostasis, revealing for the first time a central role for 3D genome architecture in regulating regenerative gene expression, and identifying cohesin as a critical regulator of nerve regeneration.

## RESULTS

### Neuronal cohesin depletion abrogates nerve regeneration

Given the role of cohesin and CTCF in 3D genome organization and chromatin looping, we took advantage of published transcriptomic datasets from purified DRG neurons at homeostasis and following nerve injury (27) as well as ChIP-seq datasets of CTCF and structural maintenance of chromosomes 1 (SMC1, cohesin subunit) from neuronal tissue (28–30) to examine the binding of CTCF and cohesin to the promoters of genes at homeostasis and after axonal injury. A high proportion of neuronal promoters displayed binding sites for cohesin at both homeostatic and injury conditions. Specifically, 47.5% of the injury-responsive promoters showed binding sites for cohesin, while only 27.3% showed binding sites for CTCF (SI Appendix, Fig. S1A). Interestingly, genes with cohesin binding sites were enriched for biological processes associated with regenerative ability including nervous system development, neuronal differentiation, axon projection and guidance, microtubule organization, axon transport, and synapse organization (13, 31, 32) (SI Appendix, Fig. S1B). However, genes with CTCF binding sites were less enriched for such biological processes (SI Appendix, Fig. S1B). By screening published transcriptomic datasets from purified cortical and retinal ganglion neurons in regenerative conditions (33, 34), we found that neuronal promoters displayed binding sites for cohesin and CTCF, and that a higher proportion of genes showed preferential binding to cohesin over CTCF, as in DRG neurons (SI Appendix, Fig. S1A). Although genes with cohesin binding sites showed a higher enrichment for biological processes associated with regenerative ability, genes with CTCF binding sites were also associated with multiple regenerative pathways (SI Appendix, Fig. S1B). Altogether, these data suggest that 3D genome architecture mechanisms might regulate gene expression that is required for axon regeneration in response to injury and that cohesin might have a preferential role with respect to CTCF.

To test this hypothesis, we assessed nerve regeneration in cohesin-depleted versus WT DRG neurons. To this end, we conditionally deleted the cohesin subunit *Rad2*1, which is required for to the integrity of the cohesin complex (35), as well as cohesin-mediated chromatin looping and domain organization (7–10). We injected AAV-Cre-GFP or AAV-GFP in the sciatic nerves of *Scc1 flox/flox* mice (35) four weeks before a sciatic nerve crush (SNC). Nerve regeneration was assessed by measuring the intensity of Stathmin-like 2 (STMN2 or SCG10) immunolabelling, as a marker of regenerating axons (36). Neuron-specific loss of RAD21 (SI Appendix, Fig. S2A-C) resulted in a strong impairment of nerve regeneration at 7 days after nerve crush (Fig. 1A-C). Furthermore, 14 days after nerve crush, we injected cholera toxin subunit B (CTB) in the *tibialis anterioris* and *gastrocnemius* muscles, to specifically trace DRG neurons that regenerated their axons across the injury site into the muscle. The number of CTB-positive DRG neurons was significantly reduced in RAD21-depleted neurons (Fig. 1D-E). Consistently, we found a severe reduction in epidermal reinnervation 18 days post injury, as assessed by counting the number of Ubiquitin C-terminal hydrolase L1 (UCHL1 or PGP9.5) positive fibres in the hind paw interdigital skin (Fig. 1F-H). This defect in nerve regeneration was not due to neuronal death, as established by staining for active-caspase 3 and by counting the number of DRG neurons per surface unit (SI Appendix, Fig. S2D-F); instead, it was associated with loss of regenerative associated genes (RAGs) activation as shown by ATF3 and c-Jun expression studies (SI Appendix, Fig. S3).

**Figure 1.**
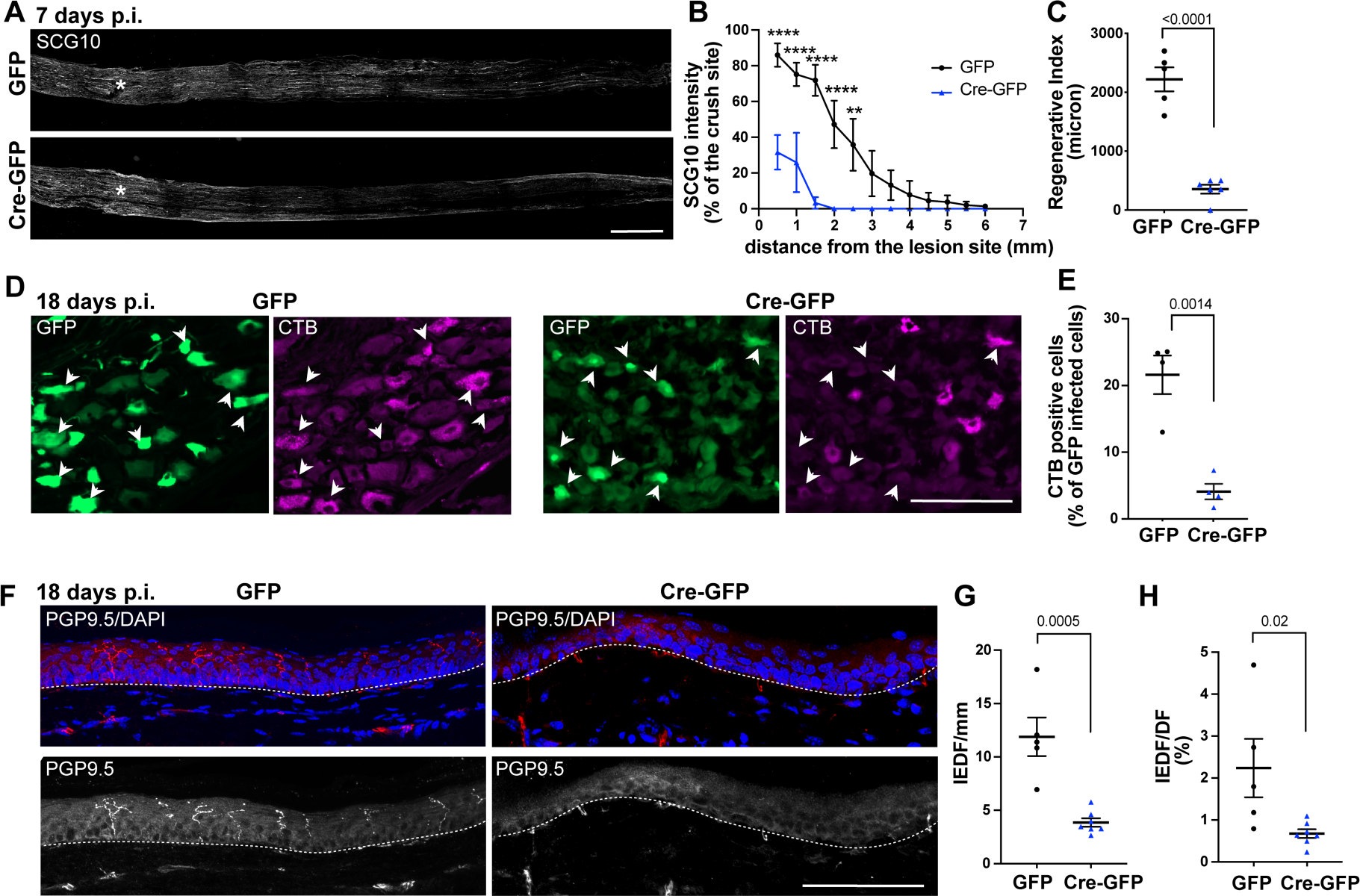
Loss of cohesin impairs nerve regeneration. **(A-B)** Representative micrographs and quantification of SCG10 intensity at the indicated distances from the lesion site in sciatic nerves from AAV-GFP or AAV-Cre-GFP injected *Scc1*flox/flox mice at 7 days following SNC. Asterisk marks the lesion site; scale bar, 1mm. (mean±s.e.m of *n*=5 nerves from 4 AAV-GFP mice and *n*=6 nerves from 4 AAV-Cre-GFP mice; **P<0.05, ****P<0.0001, 2-way Anova, Sidak’s multiple comparisons test). **(C)** Bar charts of the regeneration index in AAV-GFP or AAV-Cre-GFP injected *Scc1*flox/flox mice at 7 days after SNC (mean±s.e.m of *n*=5 nerves from 4 AAV-GFP mice and *n*=6 nerves from 4 AAV-Cre-GFP mice; two-sided unpaired Student’s *t*-test). **(D)** Micrographs showing GFP (green) and CTB (magenta) signal in DRG 4 days after injection of CTB in the tibialis anterioris and gastrocnemius muscle 14 days after SNC. Arrowheads mark GFP positive neurons. Scale bar, 100 μm. **(E)** Bar graphs of the CTB positive neurons as percentage of GFP expressing neurons (mean±s.e.m of *n*=4 mice; two-sided unpaired Student’s t-test). **(F)** Representative micrographs of PGP9.5 immunostaining with 4′,6-diamidino-2-phenylindole (DAPI) counterstaining in the interdigital hind-paw skin from AAV-GFP or AAV-Cre-GFP injected *Scc1*flox/flox mice 18 days after SNC. The dashed lines indicate the boundary between the epidermis and dermis; scale bar, 100 µm. **(G-H)** Quantification of the number of intra-epidermal fibres (IEDF) per millimeter of interdigital skin and the percentage of IEDF versus dermal fibres (DF) (mean±s.e.m of *n*=5 skins from 4 AAV-GFP mice; *n*=7 skins from 4 AAV-Cre-GFP mice; two-sided unpaired Student’s *t*-test).

Together, these data show that cohesin is required for nerve regeneration and suggest that cohesin-dependent chromatin 3D architecture mechanisms might underpin the regenerative program after injury.

### Cohesin is required for the activation of neuronal regenerative genes and pathways following injury

Next, we investigated whether cohesin is required for the establishment of the gene expression programme that follows nerve injury and supports regeneration. To this end, we performed RNA-seq from WT or *Rad21* conditionally deleted DRG neuronal nuclei in naïve conditions as well as three days after SNC versus sham control. We took advantage of the isolation of nuclei tagged in specific cell types (INTACT) mouse (37) and generated INTACT-AdvillinCre mice (see Methods) that express a super folded GFP-Myc tagged version of SUN1 (nuclear membrane protein) in Advillin positive cells following tamoxifen injection (SI Appendix, Fig. S4A). Since Advillin is expressed in all sensory neurons (38), this allows purification of the DRG sensory neuronal nuclei at high yield and purity (SI Appendix, Fig. S4B-D). We then crossed INTACT-AdvillinCre mice with *Scc1*flox/flox mice to induce uniform depletion of RAD21 in DRG neurons (SI Appendix, Fig. S5). Sixteen days after tamoxifen injection, RNA-seq was performed. Spearman correlation and principal component analysis (PCA) of the RNA-seq data showed high reproducibility between replicates (SI Appendix, Fig. S6). RNA-seq confirmed a 70% loss of *Rad21* expression in DRG neurons. RAD21 depletion induced downregulation of 711 and upregulation of 283 genes in naïve conditions (FDR<0.05) (SI Appendix, Fig. S7A). Following injury, the number of differentially expressed genes increased, with 1093 downregulated and 694 upregulated genes (FDR<0.05) (SI Appendix, Fig. S7B).

To investigate the biological significance of the genes affected by the lack of cohesin, we performed a gene ontology analysis of the differentially expressed genes in *Rad21* KO versus WT neurons in different conditions. In naive condition, *Rad21* deletion induced the downregulation of genes involved in neuronal-specific functions, such as ion transport and neurotransmitter signalling (Fig. 2A, first column). In agreement with previous findings (15, 27, 31, 32, 39, 40), in WT neurons nerve injury triggered regenerative pathways as shown by the expression of genes involved in transcription, immune regulation, nervous system developmental processes, signal transduction, cytoskeleton remodelling, circadian rhythms, axonogenesis, angiogenesis, cell adhesion, reactive oxygen species signalling, and downregulation of genes involved in ion transport and synaptic transmission (Fig. 2A, second column, and SI Appendix, Fig. S7C). However, in RAD21 depleted neurons most of the genes belonging to regenerative pathways either failed to be activated at all by injury, were induced at lower levels, or were downregulated compared to WT neurons (Fig. 2A, third and fourth columns). Genes that were downregulated after injury in WT neurons were also downregulated in *Rad21* KO neurons. Genes that remained inducible or were upregulated in the absence of cohesin were enriched for categories involved in the inflammatory response (Fig. 2A, third and forth columns). This suggests that cohesin is specifically required for the activation of the regenerative transcriptional program in response to injury.

**Figure 2.**
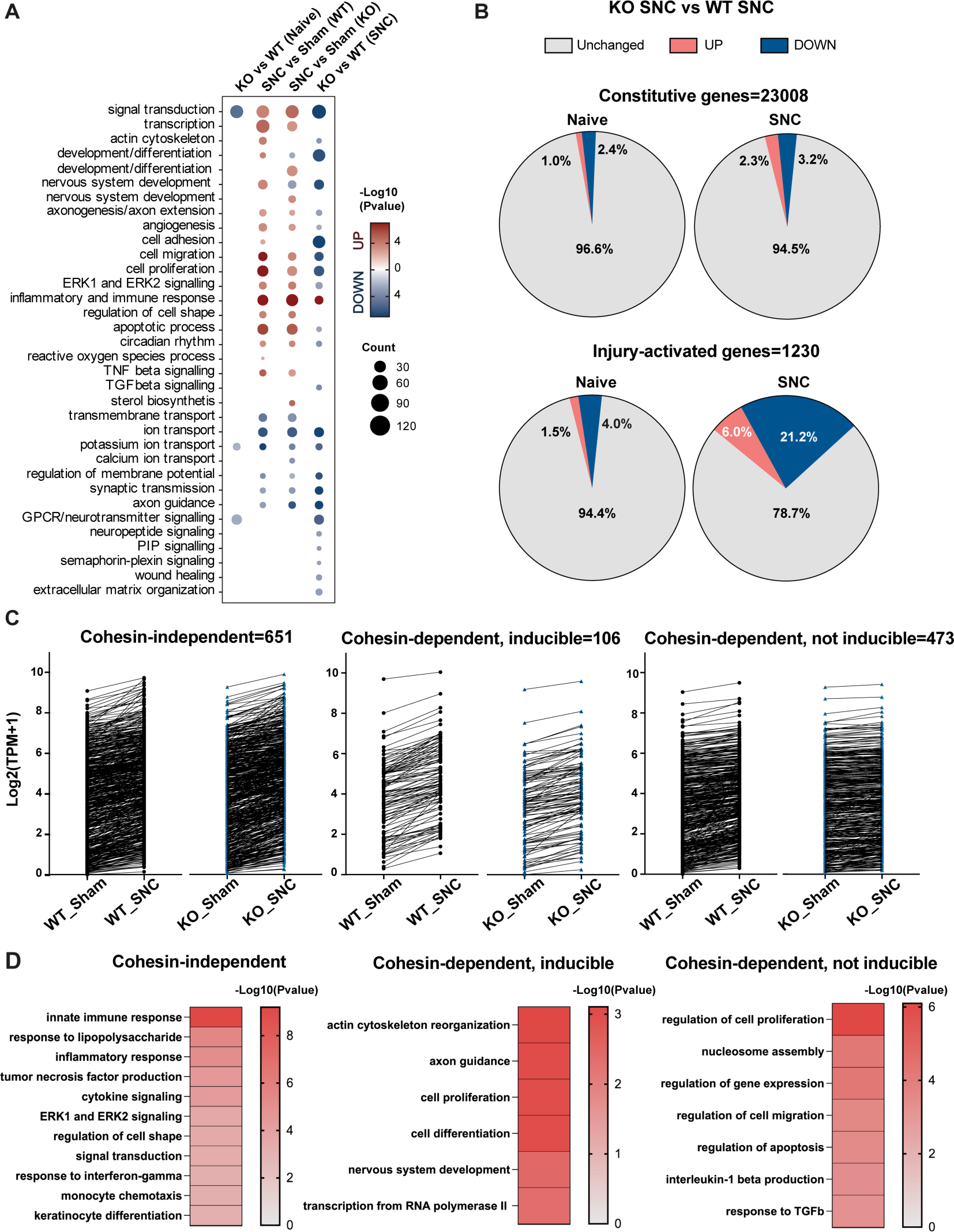
Cohesin-dependent and independent genes. **(A)** Dot plot of the semantically clustered gene ontology (GO) biological process categories of the upregulated (red) and downregulated (blue) genes in the indicated conditions. Color code reflects the P-value (modified Fisher’s exact P ≤ 0.005) and the size of the dot the gene count in each category (gene count>6). **(B)** Pie charts of the expression of constitutive (top) and injury-activated (bottom) genes in *Rad21* KO neurons in naïve and injured conditions (*n*=3 independent samples; FDR<0.05). **(C)** Line plots of the expression in the indicated conditions of the cohesin-independent and dependent (inducible and non-inducible) genes (TPM=transcripts per million). **(D)** Heatmaps of the semantically clustered GO biological process categories of the cohesin-dependent and independent genes. Color code reflects the P-value (modified Fisher’s exact P ≤ 0.001).

To directly explore this hypothesis, we analysed constitutive homeostatic genes (genes that were not differentially regulated after SNC versus Sham in WT neurons, n=23008) and genes that were activated by injury (upregulated after SNC vs Sham) in WT neurons (n=1230), separately. Only between 3.4 and 5.5% of the constitutive genes were differentially expressed in *Rad21* KO versus WT neurons in naïve and injury conditions, respectively (Fig. 2B, top). In striking contrast, 21.2% of the injury-activated genes were downregulated in *Rad21* KO versus WT neurons following injury, but not in naïve conditions (Fig. 2B, bottom). Only 6.0% of the injury-activated genes were upregulated in *Rad21* KO neurons following injury.

We then assessed the expression pattern of the 1230 injury-activated genes, in *Rad21* KO and WT neurons. We found that 651 genes were still fully inducible and upregulated in *Rad21* KO neurons after SNC versus Sham to WT levels or higher (Fig. 2C, and SI Appendix, Fig. S7D, cohesin-independent). These cohesin-independent genes were enriched for immune-related functions (Fig. 2D, left). We found 579 genes that were either downregulated in injured cohesin depleted neurons with respect to WT or that were not inducible after injury. Among these 579 cohesin-dependent genes, 106 were partially inducible but exhibited a lower baseline expression and a decreased response to injury (Fig. 2C, and SI Appendix, Fig. S7D, cohesin-dependent, inducible) and 473 genes failed to respond to injury (Fig. 2C, and SI Appendix, Fig. S7D, cohesin-dependent, not inducible). Cohesin-dependent genes were mainly enriched for actin cytoskeleton remodelling, axon guidance, neuron development, signalling, RNA polymerase II dependent transcription, nucleosome assembly, and immune-modulation pathways (Fig. 2D, middle and right). Accordingly, we found that the lack of cohesin impaired the injury-driven activation of RNA polymerase II, with only modest effects on the steady-state levels of the active enzyme (SI Appendix, Fig. S8).

Altogether, these data indicate that cohesin is required for the transcriptional response to injury that is related to the regenerative potential by controlling the expression of genes involved in regenerative pathways.

### Cohesin-dependent 3D chromatin architecture is required for the activation of neuronal regenerative genes following injury

To gain insights into the role of the 3D genomic architecture of DRG neurons in the transcriptional regulation at homeostasis versus nerve injury, we next performed Hi-C from WT and RAD21 depleted DRG neurons three days following SNC versus sham by using the same inducible-INTACT-AdvillinCre mouse model used for RNA-seq. Stratum-adjusted correlation coefficients (SCC) scores of the Hi-C data showed high reproducibility between replicates (SI Appendix, Fig. S9). After aligning and filtering an average of 712 million Hi-C sequenced read pairs, we obtained an average of 210 million long-range (> 20 Kb) intra-chromosomal contacts (MAPQ>0). A/B compartments and chromatin 3D domains were called on replicate-pooled, balanced Hi-C contact matrices (see SI Appendix, Supplementary Methods). We identified 7662, 7683, and 7753 domains in WT neurons in naïve, sham, and injury conditions, respectively.

Hi-C revealed that *Rad21* KO in DRG neurons resulted in mild effects on global genome A/B compartmentalization (SI Appendix, Fig. S10), whereas it strongly disrupted the organization of 3D chromatin domains in both naïve and injury conditions (Fig. 3A-D). Cohesin-depleted neurons showed a loss of insulation at 3D domain boundaries (Fig. 3A and C) and led to the loss of 5,873 and 4,598 domains in naïve and injury conditions, respectively. We also observed a genome-wide reduction in contact frequency as a function of the genomic distance in cohesin-depleted vs WT neurons (Fig. 3C). The distance between the “within domains” and “between domains” curves was strongly decreased indicating a loss of the chromatin contacts in cohesin-depleted neurons (Fig. 3B). We then normalized and balanced the Hi-C contact frequencies (see SI Appendix, Supplementary Methods) coupled with RNA-seq to identify changes in chromatin interactions within domains and their correlation with gene expression between WT and *Rad21* KO neurons. We identified 6,624 and 5,359 domains with a lower interaction frequency (FDR<0.05) in naïve and injury conditions, respectively, (Fig. 3D). Differentially expressed, mainly downregulated, genes were more strongly associated with domains showing a decreased frequency of interaction (Fig. 3E).

**Figure 3.**
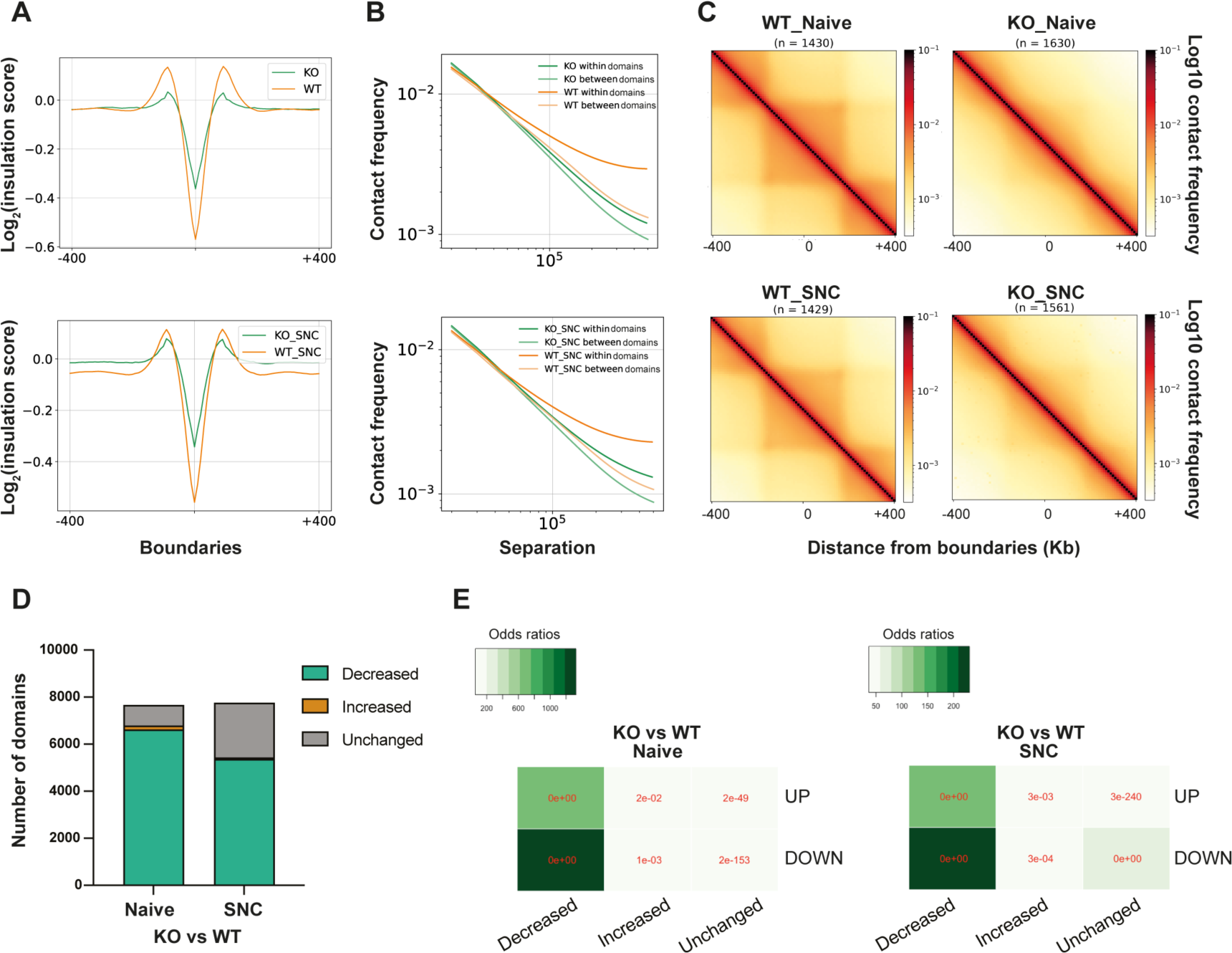
Loss of cohesin leads to a decrease in chromatin interactions. **(A)** Genome-wide plot of the averaged insulation scores around strong boundaries (the size of flanks around each strong boundary is 400 Kb) in the indicated conditions. **(B)** P(s) curve (contact probability vs genomic distance) within and between genomic domains of length 300-500 Kb for the indicated conditions. **(C)** Average Hi-C contact matrices at 10-Kb resolution of the indicated number of 3D chromatin domains of length 300-400 Kb in the indicated conditions. **(D)** Bar charts of the number of 3D chromatin domains showing an average increased, decreased, and unchanged frequency of contacts (*n*=3 independent samples; FDR<0.05). **(E)** Odds ratios analysis of the association between up- and down-regulated genes and differentially expressed genes present in increased, decreased, or unchanged domains; the numbers in red represent the P-value given by two-sided Fisher’s exact test.

Downregulated genes residing in genomic domains that were lost or showed a reduction in strength in *Rad21* KO neurons were enriched for regenerative pathways, such as axon extension and guidance, nervous system development, neuronal differentiation, circadian rhythm, angiogenesis, actin cytoskeleton remodelling (15, 31, 32, 39, 40) (Fig. 4A). The previously identified 579 cohesin-dependent genes, including several RAGs, were preferentially found within domains that were lost or showed a reduction in strength (Fig. 4B-D). Altogether, these data indicate that cohesin activates the transcriptional response to injury that is related to the regenerative potential by controlling the 3D chromatin architecture of genes involved in regenerative pathways (Fig. 4E).

**Figure 4.**
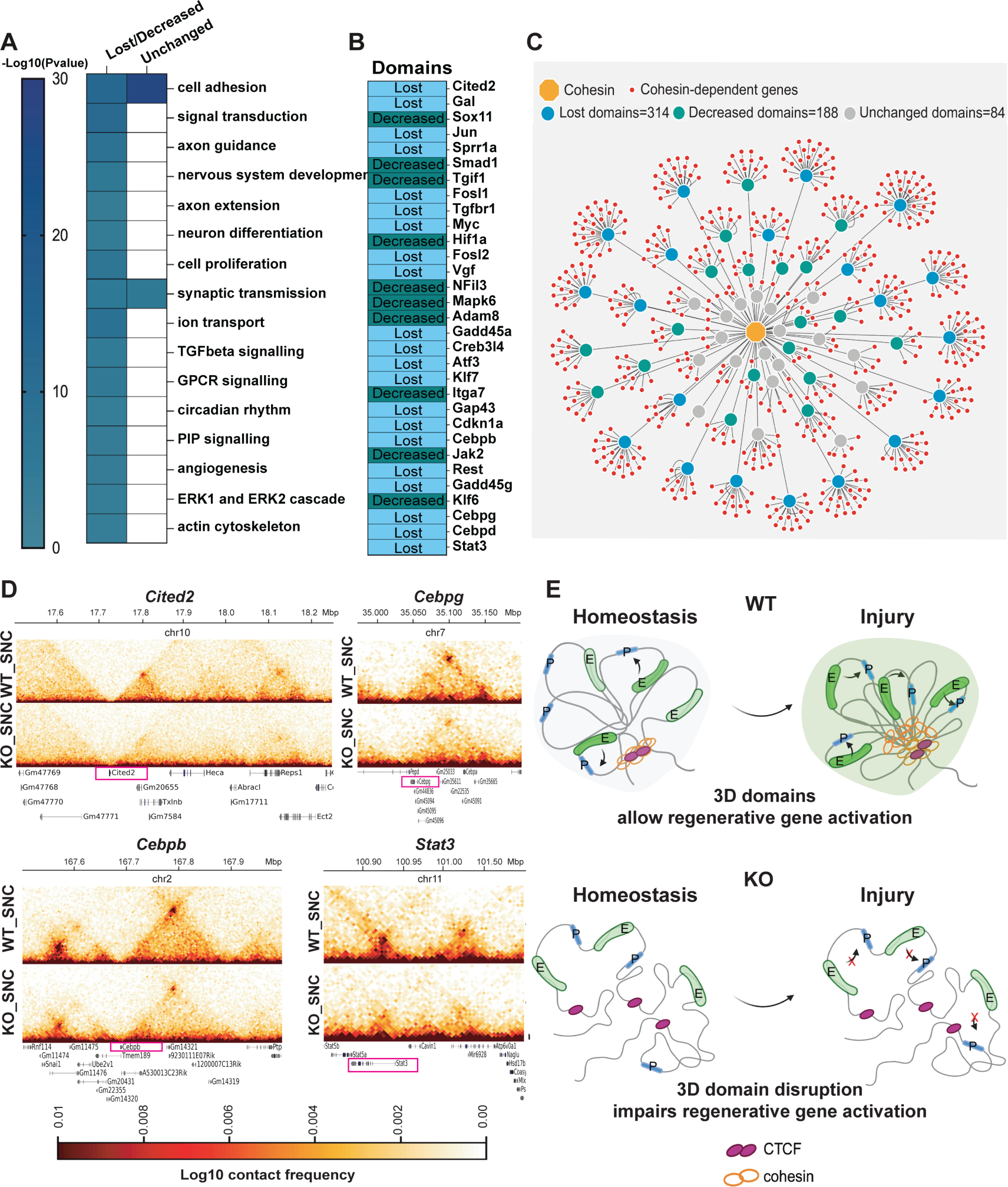
Regenerative genes reside within cohesin-dependent chromatin domains. (**A**) Heatmap of the semantically clustered gene ontology (GO) biological process categories of the downregulated genes in *Rad21* KO injured neurons residing within domains that were either lost or showed a decreased contact frequency and unchanged domains. Color code reflects the P value (modified Fisher’s exact P ≤ 0.001). **(B)** Heatmap showing the residence of RAGs within genomic domains that were either lost or showed a decreased interaction frequency in *Rad21* KO neurons. **(C)** Network visualization of the injury-activated, cohesin-dependent genes (red) (cohesin is depicted in orange). Cohesin-dependent genes are preferentially associated with lost domains (blue=314) and domains with decreased frequency (green=188) with respect to unchanged domains (gray=84). Edges connecting the genes to their genomic domains define the fold change of Hi-C interactions in SNC_KO vs SNC_WT. For a better visualization, all the domains on each chromosome are visualized as a unique circle. **(D)** Example of Hi-C maps of the contact frequency in the indicated conditions within 3D genomic domains. The contacts between the three successive bins (the one containing TSS and its two neighbors) and bins within a genomic distance of 500 Kb are extracted from 5-Kb Hi-C contact matrices. **(E)** Cohesin facilitates the formation of 3D genomic domains where regenerative genes are co-regulated. Loss of cohesin disrupts the architecture of 3D genomic domains impairing the activation of the regenerative program.

In cortical neurons, cohesin is required for the activation of activity-dependent genes that form long-range chromatin loops (41). To explore whether cohesin facilitates long-range chromatin loops between the promoters of injury-responsive genes and their regulatory elements, we leveraged recently published deep Hi-C studies in mouse cortical neurons (42). We found that injury-activated genes were involved in longer loops compared to constitutive homeostatic genes (Fig. 5A). Importantly, cohesin-dependent genes were associated with more frequent and longer loops compared to cohesin-independent genes (Fig. 5B-D). Several RAGs were found among the injury-activated genes and involved in longer loops (Fig. 5D). These data suggest a key role for cohesin-dependent 3D genome organisation in the coordination of the transcriptional response to injury in DRG neurons, by facilitating long-range chromatin loops at regenerative genes.

**Figure 5.**
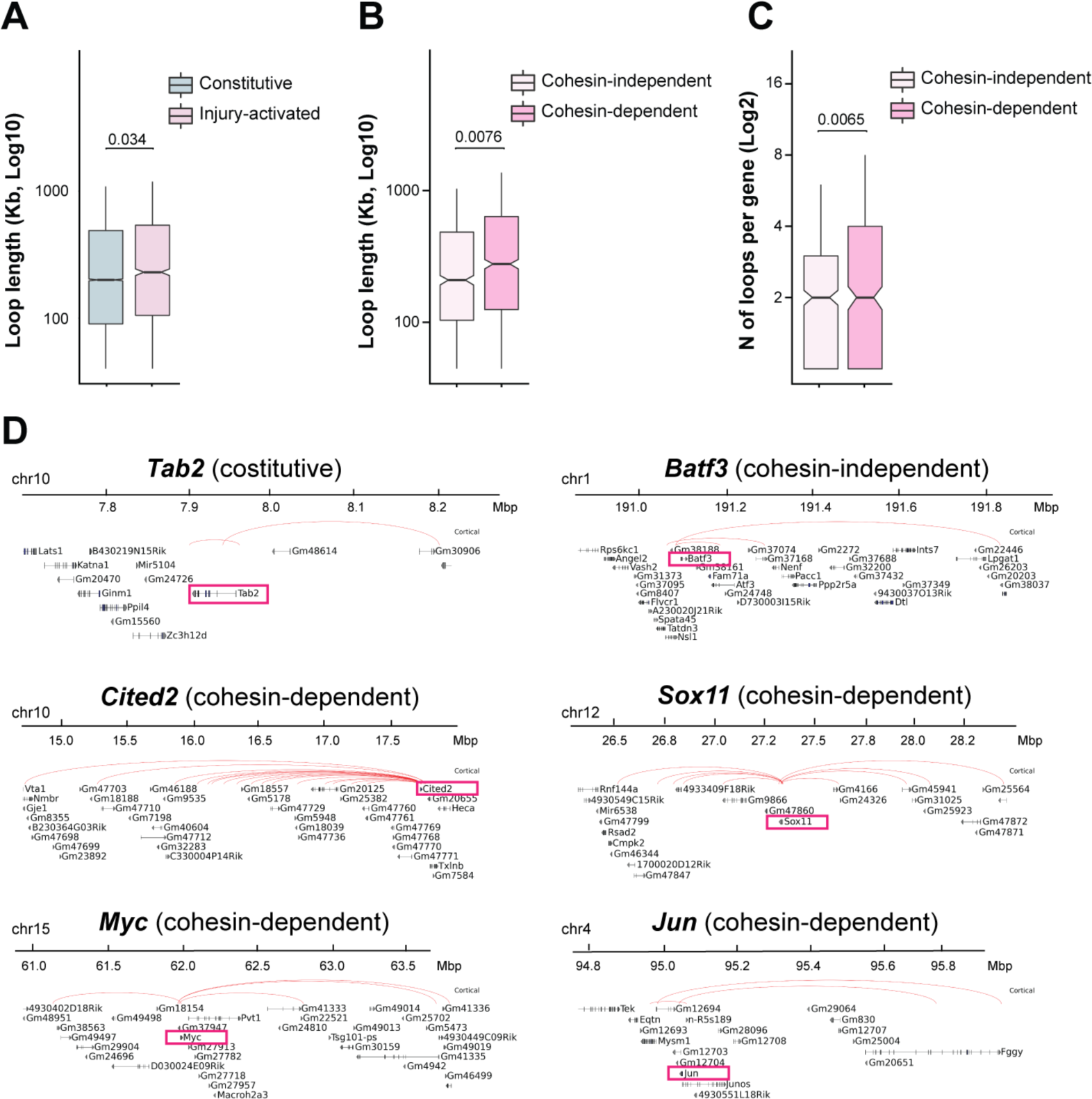
Cohesin-dependent genes engage in longer and more frequent chromatin loops in cortical neurons. **(A)** Box plot of the chromatin loop length for injury-activated and constitutive genes in cortical neurons. P-values are computed from a two-sided unpaired Student’s *t*-test. **(B)** Box plot of the chromatin loop length for cohesin-dependent and independent genes in cortical neurons. P-values are computed from a two-sided unpaired Student’s *t*-test. **(C)** Box plot of the number of chromatin loops for cohesin-dependent and independent genes in cortical neurons. P-values are computed from a two-sided unpaired Student’s *t*-test. **(D)** Examples of loops at constitutive, cohesin-dependent and independent genes.

**Figure 6.**
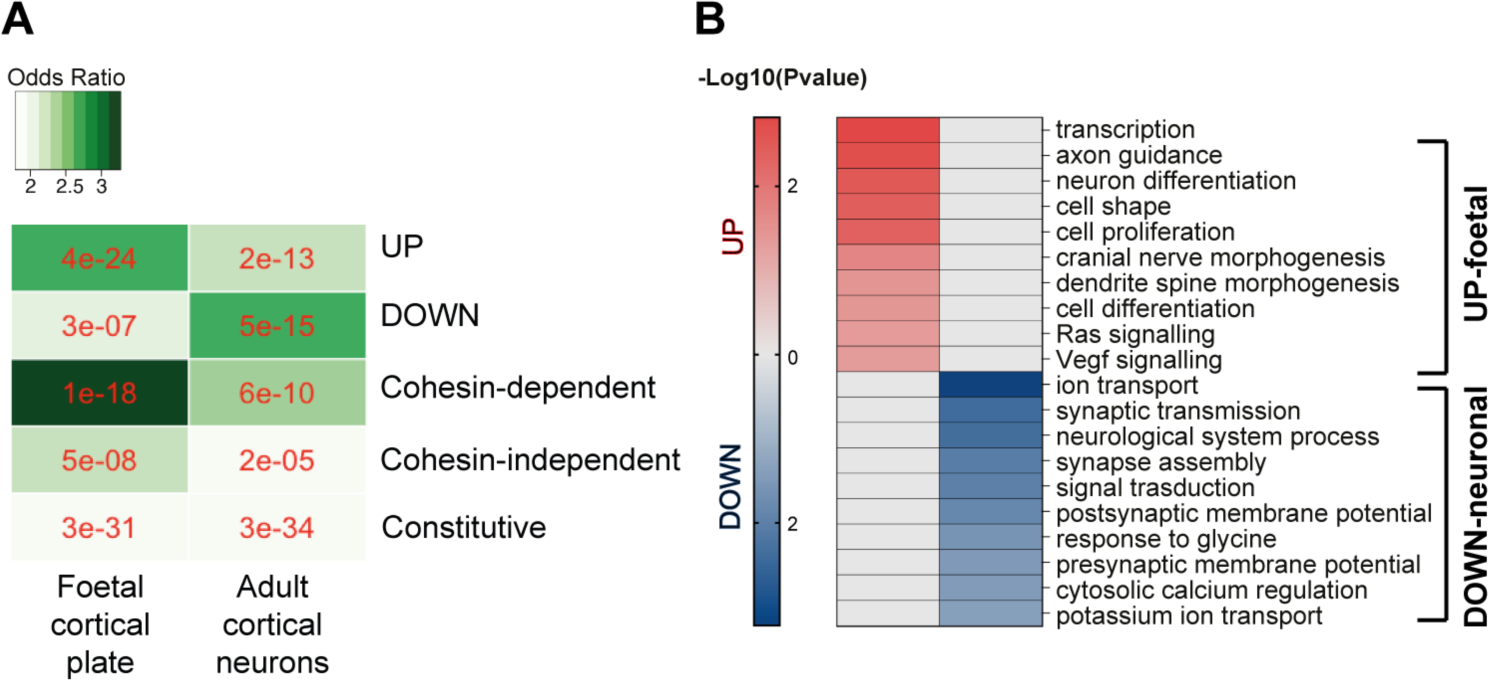
Injury-responsive genes display a more immature chromatin looping signature after injury. **(A)** Odds ratios analysis of the association between up-, down-regulated genes, cohesin-dependent and independent genes, constitutive genes, versus genes associated to foetal cortical plate and adult neuron E-P loops; the numbers in red represent the P-value given by two-sided Fisher’s exact test. **(B)** Heatmap of the semantically clustered gene ontology (GO) biological process categories of the upregulated genes associated with foetal cortical plate and downregulated genes associated with adult neuron E-P loops. Color code reflects the P-value (modified Fisher’s exact P ≤ 0.001).

### Transcriptional reprogramming after injury is associated with a more immature chromatin looping signature

Neuronal regeneration has been associated with a regression towards a transcriptional signature that mimics a less mature developmental state (33, 39). The 3D chromatin architecture changes during neuronal development, with the formation of chromatin loops and domains correlated with neuronal gene activation (42–44). To explore whether the injury-induced “transcriptional dematuration” involves a change in chromatin looping towards a state resembling a less mature development stage, we leveraged a Hi-C study on nuclei isolated from 18-24 weeks post-conception human foetal cortical plate (the period when neurons are transitioning towards functional maturation) and neurons from adult prefrontal cortex (44). We found that downregulated genes after injury preferentially overlapped with genes that were associated with E-P loops that were more specific to adult neurons. Conversely, upregulated genes after injury, and in particular cohesin-dependent genes, preferentially overlapped with genes that were associated with E-P loops that were more specific to foetal cortical plate (Fig. 5A). Constitutive genes did not show any preferential overlap between foetal cortical plate versus adult neurons (Fig. 5A). Together, these data suggest that injured neurons assume a transcriptional signature associated to a chromatin looping profile similar to that of immature, developing neurons. Downregulated genes associated with adult human neurons E-P loops were enriched for mature neuron functions, such as ion transport and synaptic transmission; upregulated genes associated with foetal E-P loops were enriched for genes related to transcription, axon guidance, dendrite morphogenesis, and Ras and VEFG signaling (Fig. 5B).

These data suggest that the “transcriptional dematuration” in response to injury is associated with a rewiring of chromatin looping towards a 3D chromatin architecture characteristic of a less mature neuronal developmental stage.

## DISCUSSION

Our work provides a map of the 3D chromatin architecture of adult mature DRG sensory neurons *in vivo* at homeostasis and after injury. It showed that the 3D chromatin structure of adult mature neurons is important for the regenerative response to injury.

We initially observed that cohesin binding sites are present in a significantly high percentage of neuronal injury-responsive genes. Cohesin generates chromatin loops that facilitate long-range chromatin contacts between gene promoters and their enhancers within 3D genomic domains, and, in collaboration with CTCF, participates in the insulation of such domains. (9, 45). Others have shown that co-expression of CTCF with other TFs contributes to promoting axon growth (46). While we also found that injury-responsive genes were enriched for binding sites for CCCTC-binding factor (CTCF), these were significantly less numerous than the ones identified for cohesin. Additionally, while we observed a modest impairment in nerve regeneration after neuronal conditional deletion of CTCF (15), disruption of the cohesin complex via *Rad21* deletion in sensory neurons abrogated nerve regeneration. This prompted us to investigate the role of 3D chromatin folding and cohesin at homeostasis and after nerve injury.

Here, we show that the integrity of 3D chromatin organization is critical for nerve regeneration. In accordance with previous findings, we observed that cohesin depletion led to loss of 3D genomic domains, without affecting genome compartmentalization (8, 9). Such extensive alteration of 3D genome architecture resulted in the downregulation of genes that support neuronal function, including ion channels, synaptic transmission, and cell adhesion in naïve neurons, similar to what has been found in cortical neurons (41), suggesting that this is a conserved mechanism, supporting its physiological relevance in neuronal homeostasis. However, the most severe changes in gene expression were found after injury, where we observed a further downregulation of neuronal-specific genes and of several transcripts that are normally activated in response to nerve injury in WT neurons. Interestingly, it was recently demonstrated that cohesin is required for gene activation following stimulus rather than constitutive gene expression, for example for the induction of activity-dependent genes in cortical neurons and inflammatory genes in macrophages (41, 47).

Lack of cohesin did not affect all the genes that are activated in response to injury. While genes that are involved in immune and inflammatory functions were independent from cohesin for their activation, genes affected by cohesin depletion were enriched for biological pathways that have been reported to be pro-regenerative (15, 31, 32, 39, 40). These included a high proportion of RAGs (13, 14), which were found enriched within chromatin contacts that were disrupted by cohesin depletion. The downregulation of neuron-specific genes has been linked to a neuronal de-maturation process that facilitates axon growth (33, 39). These data suggest that downregulation of neuron maturation genes might not be sufficient to promote axon growth if it is not followed by activation of the full regenerative program.

Cohesin-dependent genes, in particular the ones that lost their inducibility after injury in *Rad21* KO neurons, were involved in longer loops with respect to cohesin-independent ones in cortical neurons. This suggests that 3D chromatin architecture after nerve injury might be required to facilitate enhancer-promoter contacts at regenerative genes. It has been shown that neuronal maturation is associated with an increased frequency of E-P loops that regulate neuronal activity related genes (44). We found that the downregulated genes after nerve injury were associated with E-P loops specific to adult human neurons. We also found that nerve injury induces upregulation of genes related to axon guidance, dendrite morphogenesis, signaling cascades, and regulation of transcription that were associated with E-P loops specific to foetal cortical plate. Interestingly, many genes that are upregulated in response to injury are involved in the regulation of transcription and have been described to be kept repressed via repressor-loops in mature human neurons at homeostasis (44). This suggests that, in addition to the loss of chromatin accessibility (48), the rewiring of chromatin looping across development might underpin the poor regenerative capacity of adult neurons and that this phenomenon seems to be conserved between mice and humans. These findings are also in line with loss of function mutations in the cohesin complex that are associated with neurodevelopmental diseases (49–51).

In summary, this work overcomes the limitations of previous multiomic studies performed in bulk DRG tissue (15), highlights a previously uncharacterized role for chromatin 3D architecture in controlling the nerve regenerative response after injury, and provides a 3D chromatin map of sensory neurons in their physiological and injury state. Along with chromatin accessibility and transcription factor activity and binding, 3D chromatin mechanisms, such as promoter-enhancer interactions, should be considered as an added layer of transcriptional regulation in the response to nerve injury. Furthermore, the presence of cohesin binding sites in a high percentage of genes activated in central neurons in conditions where their regeneration capacity has been stimulated, suggests that 3D chromatin mechanisms might also play a role in the central nervous system injury-response, opening new avenues for repair.

## METHODS

### Mice

Scc1flox/flox mice (gift from Dr. Matthias Merkenschlager) have been described (35). INTACT-Scc1flox/flox-AdvillinCre and INTACT-AdvillinCre mice were generated by crossbreeding INTACT mice (37) (129-Gt(ROSA)26Sor^tm5(CAG-Sun1/sfGFP)Nat^/J, JAX stock #021039) with Scc1flox/flox mice and with AdvillinCre (38) (Tg(Avil-icre/ERT2)AJwo/J, JAX stock #032027).

### Animal procedures

Animal work was carried out in accordance with the regulations of the UK Home Office. Mice were maintained in-house under standard housing conditions (12-h light/dark cycles) with 24-h access to water and a standard chow diet at temperatures between 20 °C and 24 °C and relative humidity between 45% and 65%, according to the UK Home Office Code of Practice. Males and females of 10-20 weeks were used, and animals were matched by sex and age in all experiments, whenever appropriate. INTACT-Scc1flox/flox-AdvillinCre and INTACT-AdvillinCre mice were injected intraperitoneally with tamoxifen in corn oil (100 mg/kg body weight, Sigma) once per day for 4 consecutive days, and used 16 days following the last injection, unless otherwise stated. Sciatic nerve crush (SNC) and nerve injection were performed as previously reported (15). Briefly, mice were anesthetized with isofluorane (3% induction, 2% maintenance), and buprenorphine (0.1mg/kg)/carprophen (5mg/kg) was administered preoperatively as analgesic. The sciatic nerve was exposed by blunt dissection of the *biceps femoris* and the *gluteus superficialis* and was crushed by applying pressure for 30sec with haemostatic forceps (FST). The crush site was marked by carbon powder. In control mice (sham) the sciatic nerve was exposed without crush. For nerve injection of AAV1-CAG-Cre-GFP or AAV1-CAG-GFP (Tebu-bio, CAG=cytomegalovirus (CMV) enhancer fused to the chicken beta-actin promoter), 3 μl of viral particles were injected into the nerve with a Hamilton syringe slowly, and mice were used after 4 weeks to allow complete transgene expression. All (100%) infected cells were neurons. In selected experiments, 14 days after SNC, 5 μl of 10% CTB were injected in the *gastrocnemius* and *tibialis anterior* and sacrificed 4 days afterward to trace regenerating neurons in the DRG, as a measure of muscle reinnervation.

### Replication

All attempts of replication were successful.

### Blinding

All assessments were performed in blind by two independent experimenters.

Detailed descriptions of experimental methodologies used in the study are provided in SI Appendix, Supplementary Methods.

## Supporting information

Supplementary index

## ACKNOWLEDGEMENTS

We thank the LMS/NIHR Imperial Biomedical Research Centre Flow Cytometry Facility for nuclear sorting. We thank Dr. Maria Teresa Cencioni at Imperial College London with assistance in flow cytometry analysis of sorted nuclear fractions. This work was supported by The Rosetrees Trust (SDG), the MRC (SDG), Wings For Life (SDG), the National Institute for Health Research (NIHR) Imperial Biomedical Research Centre (SDG), the National Institute of General Medical Sciences grant (1R35GM137974 to ZW), and OSU start-up (PG100125 to IP). The views expressed are those of the author(s) and not necessarily those of the NHS, the NIHR, or the Department of Health.

## AUTHOR CONTRIBUTIONS

I.P. conceived the idea, designed and performed experiments, analyzed and interpreted data, wrote the manuscript, and provided funding.

T.L. co-designed the computational experiments, co-interpreted the analysis results, analyzed the data, and wrote the manuscript.

W.G., R.T.R., and S.G. performed experiments and analyzed data.

L.Z., J.C., F.M., and G.K. performed experiments.

M.M. analyzed and interpreted data and edited the manuscript.

J.WD. K., E.A., B.C., A.C., and E.D.V. analyzed the data.

Y.Y. managed mouse colonies.

F.D.V. injected drugs and collected tissue.

Z.W. provided funding, supervised the computational experiments, and edited the manuscript.

S.D.G. provided funding, designed experiments, and wrote the manuscript.

