## Supplementary index for "Three-dimensional chromatin mapping of sensory neurons reveals that cohesin-dependent genomic domains are required for axonal regeneration"

**ABSTRACT**
The in vivo three-dimensional genomic architecture of adult mature neurons at homeostasis and after medically relevant perturbations such as axonal injury remains elusive. Here we address this knowledge gap by mapping the three-dimensional chromatin architecture and gene expression programme at homeostasis and after sciatic nerve injury in wild-type and cohesin-deficient mouse sensory dorsal root ganglia neurons via combinatorial Hi-C and RNA-seq. We find that cohesin is required for the full induction of the regenerative transcriptional program, by organising 3D genomic domains required for the activation of regenerative genes. Importantly, loss of cohesin results in disruption of chromatin architecture at regenerative genes and severely impaired nerve regeneration. Together, these data provide an original three-dimensional chromatin map of adult sensory neurons in vivo and demonstrate a role for cohesin-dependent chromatin interactions in neuronal regeneration.

**This PDF file includes:**
-Supplementary Methods
-Supplementary Figures
-References

**Supplementary Methods.**

### INTACT neuronal nuclei isolation

The nuclei isolation was performed on ice according to (1) with minor modifications. Sciatic DRG from INTACT-Scc1flox/flox-AdvillinCre and INTACT-AdvillinCre mice were washed in ice-cold Hanks' Balanced Salt Solution (HBSS, Invitrogen), collected by centrifugation, flash frozen, and stored at -80°C. The DRG pellet was homogenised with a pellet pestle in 100 µl of ice-cold homogenization buffer (0.25M sucrose, 25mM KCl, 5mM $MgCl_2$, 20mM Tricine-KOH pH 7.8, 0.04% BSA, 0.15mM spermine, 0.5mM spermidine, 60 u/ml SUPERase RNase inhibitor (Invitrogen), 5 µg/ml actinomycin D (Sigma), and EDTA-free protease inhibitors (Merck)). A 10% IGEPAL-630 solution was added to the final concentration of 0.2%, and the homogenate was further dounced with additional strokes of the pellet pestle. The nuclei sample was mixed by pipetting adding 400 µl of ice-cold homogenization buffer supplemented with 0.1 % IGEPAL-630. After letting the clumps settle, the homogenate was filtered through a 70 µm mini strainer and pre-cleared by incubating for 10 minutes with 20 µL of Protein G Dynabeads (Life Technologies). After removing the beads with a magnet, the sample was incubated for 1 hour with 60 µL of Dynabeads previously bound to 10 µL of 0.2 mg/mL rabbit monoclonal anti-GFP antibody (G10362, Life Technologies). Bead-bound nuclei were washed with 2 x 500 µl, 1 x 250 µl, and 1 x 100 µl of homogenization buffer supplemented with 0.1 % IGEPAL-630. To check specificity, aliquots of input, unbound, and bound fractions were observed at the microscope or analyzed using an Aurora flow cytometer (Cytek) following DAPI staining.

### RNA-Seq

RNA-Seq was performed from sciatic DRG neuronal nuclei isolated with the INTACT procedure from WT (INTACT-AdvillinCre) and *Rad21* KO (INTACT-Scc1flox/flox-AdvillinCre) mice in naïve conditions and 3 days after nerve crush or sham (3 biological replicates, 4 mice/replicate). INTACT beads-bound neuronal nuclei were directly resuspended in RLT buffer and RNA was extracted with RNeasy Micro kit (Qiagen) and DNase on column digestion following manufacturer's guidelines. RNA quality was assessed with Agilent 2100 Bioanalyzer (Agilent), and libraries were generated at Source Bioscience (Nottingham, UK) using the NEBNext Ultra II Directional RNA Library Prep Kit for Illumina, according to the recommended manufacturer protocol. After ligation, the NEBNext Multiplex Oligos for Illumina (Dual Index Primers Set 1) adapter sequences were ligated to each sample to allow for sequencing multiplexing. Sequencing was performed on a NovaSeq 6000, generating 150bp pair-ended reads. An average of 85 million read pairs were generated for each sample, with a total of 1.5 billion read pairs generated.

**Sequence alignment, differential expression analysis**

The raw RNA-Seq reads were mapped to the reference genome (mm10) using HISAT2 v2.2.1 (2) with --known-splicesite-infile included. The splice sites were generated with the script "extract_splice_sites.py" provided in HISAT2. featureCounts v2.0.2 (3) was used to quantify mapped reads. The differential expression analysis was conducted with DESeq2 v1.38.3 (4).

**Hi-C**

Hi-C were performed from sciatic DRG neuronal nuclei isolated with INTACT procedure from WT (INTACT-AdvillinCre) and *Rad21* KO (INTACT-Scc1flox/flox-AdvillinCre) mice 3 days after nerve crush or sham (3 biological replicates, 5 mice/replicate), or in naïve conditions (Hi-C only, 2 biological replicates, 5 mice/replicate). INTACT beads-bound neuronal nuclei were crosslinked with 2% formaldehyde for 10 min at room temperature (r.t.) and then processed for Hi-C using ARIMA Hi-C Genomics kits (A510008). Briefly, chromatin was digested with multiple enzymes, and purified proximity-ligated DNA was sheared using Bioruptor to obtain an average size of 400 bp. After DNA size selection of 200-600 bp, biotin enrichment, and end repair and adapter ligation, Hi-C libraries were prepared starting from 30-65 ng of DNA and 10 PCR cycles, using Swift Biosciences Accel-NGS 2S Plus DNA library kit (21024) and KAPA Library amplification kit (KK2620) or Arima Library preparation module (A303010). After quality control on Bioanalyzer, Hi-C libraries were sequenced on a NovaSeq 6000, generating 150bp pair-ended reads at Source Bioscience.

**Hi-C data processing**

The raw sequencing reads of each individual replicate and pooled replicates for each condition were mapped to the mouse reference genome (mm10) with Juicer (5). The Arima restriction sites were generated with the script "generate_site_positions.py" provided in Juicer. The raw Hi-C contact matrices (MAPQ > 0) in hic format were converted to cool format (6) using “hicConvertFormat” in HiCExplorer v3.7.2 (7). The reproducibility was evaluated on raw Hi-C contact matrices by stratum-adjusted correlation coefficient (SCC) from HiCRep (8, 9) at 100-Kb resolution with the smoothing parameter set to three, maximal genomic distance set to 5Mb, and performing down-sampling. For balancing Hi-C contact matrices between any two conditions, the raw Hi-C contact matrices of each individual replicate or pooled replicates in cool format from two conditions were first jointly normalized to the smallest read count using “hicNormalize” in HiCExplorer, and then balanced using “cooler balance” (6)with default parameters, which implements iterative correction and eigenvector decomposition (ICE)(10). We refer to these normalized and balanced Hi-C contact matrices as ICE-balanced.

**A/B compartments**

The A/B compartments for any two conditions performing differential analysis were obtained using “eigs-cis” in cooltools v0.5.0 (11) on replicate-pooled, ICE-balanced Hi-C contact matrices at 25-Kb and 100-Kb resolution. The GC-content is used as the phasing track for orienting and ranking eigenvectors.

**Topologically associating domains**

The topologically associating domains for any two conditions performing differential analysis were called using insulation scores implemented in cooltools v0.5.0 (11). The matrices we used for calling TADs are replicate-pooled, ICE-balanced Hi-C contact matrices at 10-Kb and 25-Kb resolution with the window sizes set to 10 and 5, respectively. If at least one boundary of a TAD in first condition is not found in second condition, we labelled this TAD as lost in second condition. If at least one boundary of a TAD in second condition is not detected in first condition, we labelled this TAD as gained in second condition. We allow $\pm$ 1 bin mismatch when checking boundary’s presence. For each TAD, we calculated the average value of log2 fold changes over each pixel located in the TAD at 10-Kb resolution and obtained P-values by computing the Wilcoxon rank-sum statistic, which were further converted to the false discovery rates (FDR) with the Benjamini-Hochberg procedure.

**Retrotranscription and q-PCR**

cDNA was synthesized from 0.5-1 μg of total RNA using the SuperScript II Reverse Transcriptase kit (Invitrogen) with both oligodT and random hexamers. QPCR was performed using Platinum SYBR Green qPCR SuperMix-UDG w/ROX (Invitrogen) on an AriaMx Real-Time PCR System (Agilent). The thermal cycling conditions were 1 cycle at 95 °C for 5 min, followed by 45 cycles at 95 °C for 20 s, 56 °C for 20 s, and 72 °C for 30 s. Primers were as follow: mTuj1F: CCCAGCGGCAACTATGTAGG; mTuj1R: GCACATACTTGTGAGAGGAGGC; mSox10F: CTTGGGACACGGTTTTCCAC; mSox10R: TCACTTTCGTTCAGCAACCTCC; mNf200F:TCAAGTGCGACGTGACGTCG; mNf200R: AGTCGGTCCAACCTCACTCG; mGfapF: CACGAAGCTAACGACTATCGC; mGfapR: AGTGCCTCCTGGTAACTGGC.

**Gene Ontology and Pathway analysis**

GO and KEGG Pathway analysis were performed using DAVID 6.8 (https://david.ncifcrf.gov/) and setting all the expressed genes in our dataset as background.

**Great Analysis**

To find genes with binding sites for CTCF and cohesin, chromosome coordinates retrieved from (12) and (13), and (14) were input into GREAT (15). The gene regulatory region was set at -3000/+1000 bp from the TSS.

**Tissue Immunohistochemistry**

Dissected DRG, sciatic nerves, and hind paw skin were fixed in 4% PFA for 2 hours and transferred to 30% sucrose at 4ºC for 5 days. The tissue was embedded in an OCT compound (Tissue-Tek) and frozen. Nerves and DRG were cryosectioned in 10 μm sections, skin sections were 20 μm thickness. For RAD21 staining in DRG, antigen retrieval was performed by heating the slides at 98°C for 3 min in 10mM Tris /1mM EDTA (pH 9.0) containing 0.1% Tween20. After washing, sections were permeabilized and blocked in 10% normal donkey or goat serum (Sigma) and 1% Triton X-100 in PBS for 1h at r.t., followed by incubation o.n. at 4ºC with the following primary antibodies in 10% normal donkey or goat serum and 0.3% Triton X-100 in PBS: anti-RAD21 (1:1000, ab217678, Abcam), anti-GFP (1:2000, ab13970, Abcam), anti-CTB (1:1000, 703, Quadratech), anti-SCG10 (1:1000, NBP1-49461, Novusbio), anti-PGP9.5 (1:200, 14730-1-AP, Proteintech), anti-CGRP (1;100, ab81887, Abcam), anti-NF200 (1:1000, N0142, Sigma), anti-

Tuj1 (1:1000, G7120, Promega), anti-active caspase 3 (1:1000), anti-ATF3 (cs-188, Santa Cruz, 1:100), anti-Jun (1:50, 9165S, Cell signalling), anti-S5 Pol-II (ab5408, Abcam, 1:1000). Alexa Fluor conjugated donkey or goat secondary antibodies (Molecular Probes) were used, according to standard protocol. Nuclei were counterstained with Hoechst (Molecular Probes).

**Image analysis**

DRG micrographs were taken at 10 or 20X magnification with a Nikon Eclipse TE2000-U mounting an Optimos Camera (Q-imaging) with a resolution of 1920x1080, or a Zeiss Axio Observer mounting a Hamamatsu Flash 4.0 fast camera with a resolution of 1998x13419, or an SP8 Leica confocal with a resolution of 1024x1024. Nerve micrographs were taken at 10x magnification using a Zeiss Axio Observer mounting a Hamamatsu Flash 4.0 fast camera with a resolution of 1998x13419. Skin micrographs were taken at 20 or 40x magnification on an SP8 Leica confocal with a resolution of 1024x1024 or 2048x2048. Images were analysed either by counting the number of positive cells per area or measuring the staining intensity against the background or calculating the percentage of cells with positive staining.

Nerve regeneration after SNC was quantified by assessing the SCG10 intensity at various distances after the crush site (which is defined by the position along the nerve length with maximal SCG10 intensity and identified by carbon powder mark) and normalized by the intensity at the injury site. The regeneration index was calculated as the distance at which SCG10 intensity is half of that at the injury site. Some nerves were excluded from analysis when, due to the processing, the integrity of the tissue was not suitable for analysis.

**Statistical analysis**

Statistical analyses were performed with GraphPad Prism software. Values are presented as dot blots with individual data points, with mean ± s.e.m. or s.d., as indicated in figure legends. All measurements were taken from distinct samples and the same sample was never measured repeatedly. Statistical analyses were designed using the assumption of normal distribution, although that assumption was not explicitly tested. Statistical comparisons included two-sided paired or unpaired Student's t-test, 1- or 2-way ANOVA followed by Sidak', Dunnett's, or Tukey's multiple comparison post hoc tests as specified in the figure legends, and one or two-sided Fisher's exact test. P-values<0.05 were considered statistically significant. The sample size was either based upon similar previously established experimental designs or calculated using an AEEC power calculator, to estimate the number of replicates required, for a difference of 1.5 and 80% to 90% power assuming a 5% significance level.

### Supplementary Figures

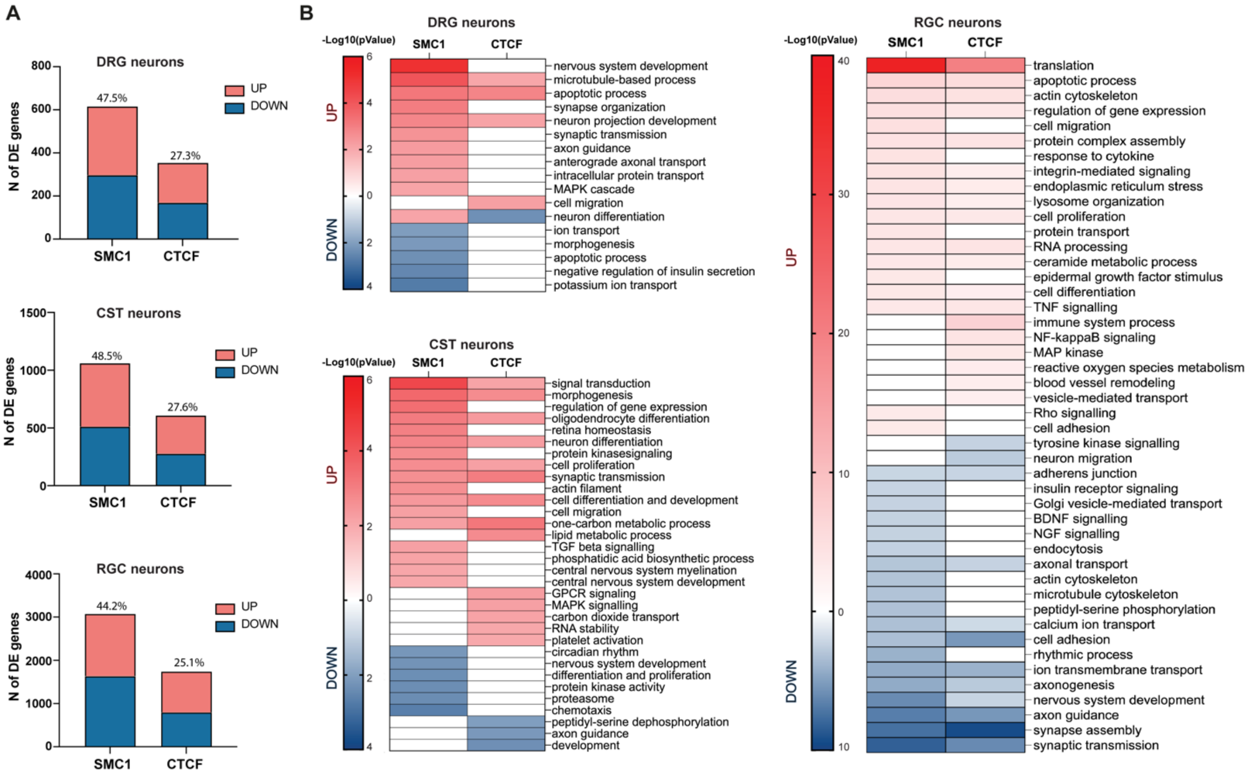


**Supplementary Figure 1. Cohesin binding sites at injury-responsive genes. (A)** Bar chart of the number of differentially expressed (DE) genes in dorsal root ganglia (DRG), cortical spinal tract (CST) and retinal ganglia cells (RGC) neurons in regenerative conditions following nerve, spinal cord, and optic nerve injury vs unjured condition (16-18), containing binding sites for the indicated proteins at the promoter level. The percentage on top of the bars represents the percentage with respect to the total number of DE genes ($n$=3 independent samples; FDR<0.05). CTCF binding sites were identified in neuronal tissue were retrieved from (12) and (13). Cohesin binding sites were identified from ChIP-seq dataset for structural maintenance of chromosomes 1 (SMC1) (cohesin subunit) (14). **(B)** Heatmap of the semantically clustered gene ontology (GO) biological process categories of the upregulated (red) and downregulated (blue) genes containing cohesin and CTCF binding sites. Colour code reflects the P-value (modified Fisher's exact $P \leq 0.001$) in each category.

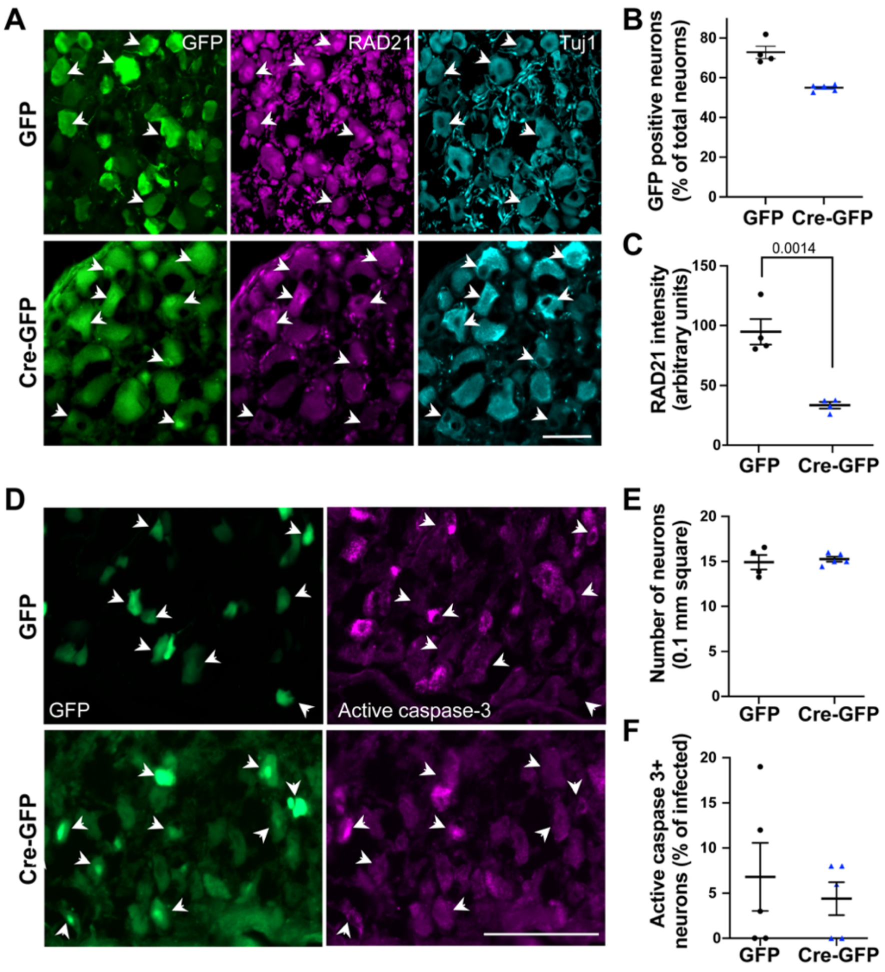


**Supplementary Figure 2. *Rad21* deletion. (A)** Micrographs showing GFP (green) and RAD21 (magenta) expression in Tuj1 (cyano) positive DRG neurons 4 weeks after AAV-GFP or AAV-Cre-GFP injection in *Scc1*flox/flox mice. Arrowheads mark GFP positive infected neurons Scale bar, 100 µm. **(B)** Scatter plot of the quantification of GFP positive infected neurons as shown in **(A)** (mean±s.e.m of *n*=4 mice). **(C)** Scatter plot of the quantification of RAD21 nuclear signal intensity as shown in **(A)** (mean±s.e.m of *n*=4 mice; two-sided unpaired Student's t-test). **(D)** Micrographs showing GFP (green) and active caspase-3 (magenta) expression in DRG from AAV-GFP or AAV-Cre-GFP injected *Scc1*flox/flox mice at 18 days after nerve crush. Arrowheads mark GFP positive infected neurons Scale bar, 100 µm. **(E)** Scatter plot of the quantification of DRG neurons per 0.1 mm square (mean±s.e.m of *n*=5 mice; two-sided unpaired Student's t-test). **(F)** Scatter plot of the quantification of active caspase-3 and GFP double positive neurons (mean±s.e.m of *n*=5 mice; two-sided unpaired Student's t-test).

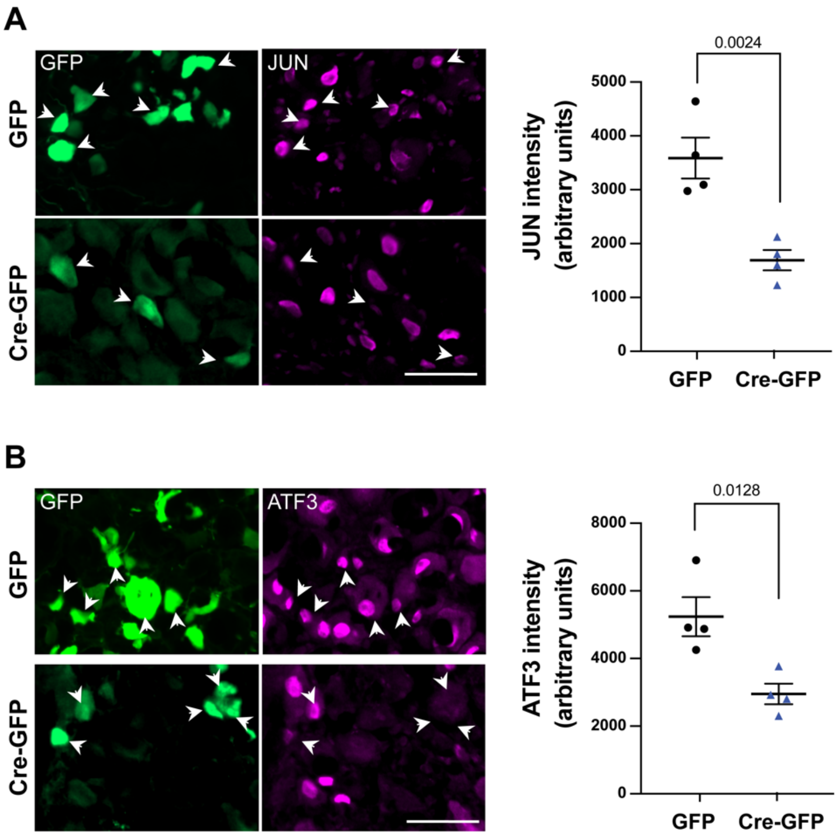


**Supplementary Figure 3. RAG expression in cohesin-depleted neurons. (A-B)** Micrographs showing GFP (green) and JUN (**A**) or ATF3 (**B**) (magenta) expression in DRG neurons 4 weeks after AAV-GFP or AAV-Cre-GFP injection in *Scc1* flox/flox mice, 7 days post injury. Arrowheads mark GFP positive infected neurons. Scale bar, 100 µm. Scatter plot of the quantification of JUN and ATF3 nuclear signal intensity (mean±s.e.m of *n*=4 mice; two-sided unpaired Student's t-test).

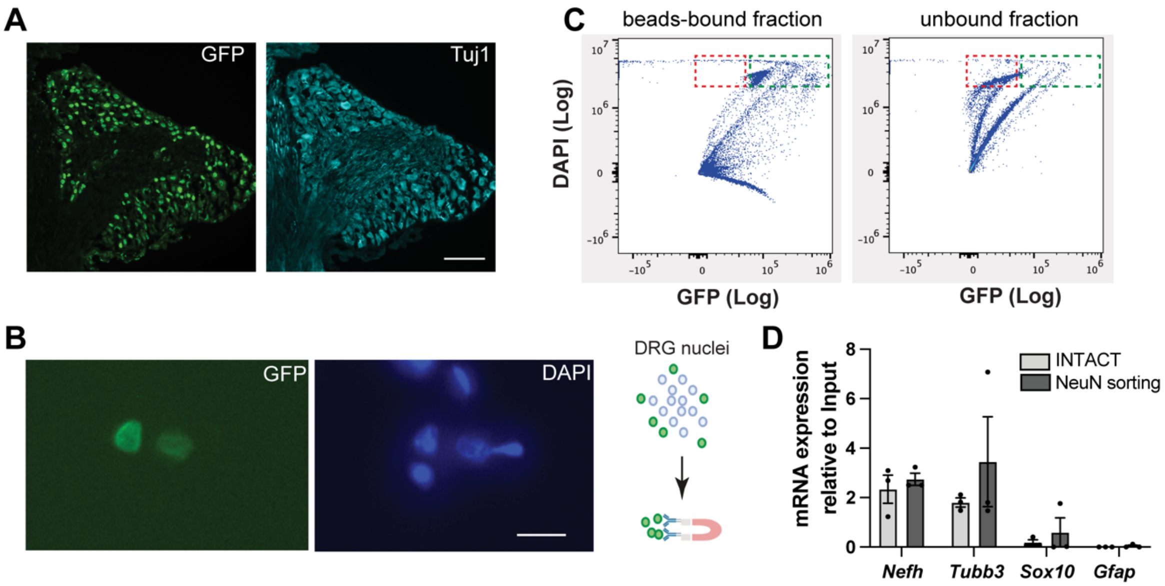


**Supplementary Figure 4. INTACT strategy. (A)** Micrographs showing SUN1-GFP (green) expression in Tuj1 (cyano) positive DRG neurons in INTACT AdvillinCre mice. Scale bar, 1 mm. **(B)** Left: micrographs showing DRG nuclei stained with DAPI before INTACT protocol purification. Scale bar, 25 µm. Right: schematic of the INTACT procedure. **(C)** Flow cytometry analysis of the beads-bound and the unbound fraction after INTACT purification. Green and red boxes show the GFP positive nuclei in the beads-bound fraction and the GFP negative nuclei in the unbound fraction, respectively. Beads-bound fraction contains only GFP positive nuclei, while some GFP positive nuclei are lost in the unbound fraction. **(D)** qPCR analysis of the expression of neuronal (*Nfh* and *Tubb3*) and non-neuronal markers (*Sox10* and *Gfap*) in the beads-bound fraction with respect to the input shows the purity of the INTACT preparation. Comparable purity was obtained by NeuN-A488 nuclear sorting.

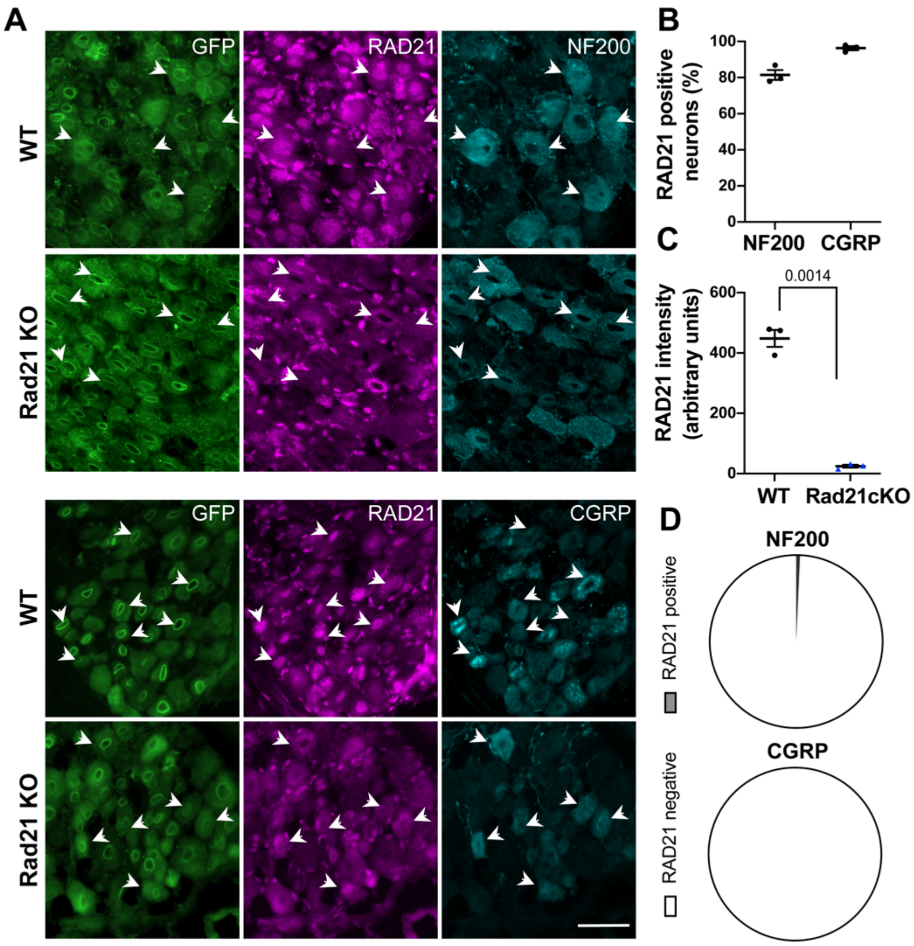


**Supplementary Figure 5. *Rad21* genetic deletion. (A)** Micrographs showing nuclear GFP (green) and RAD21 (magenta) expression in neurofilament 200 (NF200) and calcitonin G related peptide (CGRP) (cyano) positive DRG neurons after 4 consecutive daily tamoxifen injections in INTACT-AdvCre (WT) and INTACT-AdvCre-*Scc1*flox/flox (*RAD21* KO) mice. Arrowheads mark nuclear GFP positive neurons. Scale bar, 100 µm. **(B)** Scatter plot of the percentage of RAD21 and NF200 or CGRP double positive neurons in WT mice (mean±s.e.m of *n*=3 mice). **(C)** Scatter plot of the quantification of RAD21 nuclear signal intensity as shown in **(A)** (mean±s.e.m of *n*=3 mice; two-sided unpaired Student's t-test). **(D)** Pie chart of the percentage of NF200 or CGRP neurons showing RAD21 downregulation in *Rad21* KO mice (mean of *n*=3 mice).

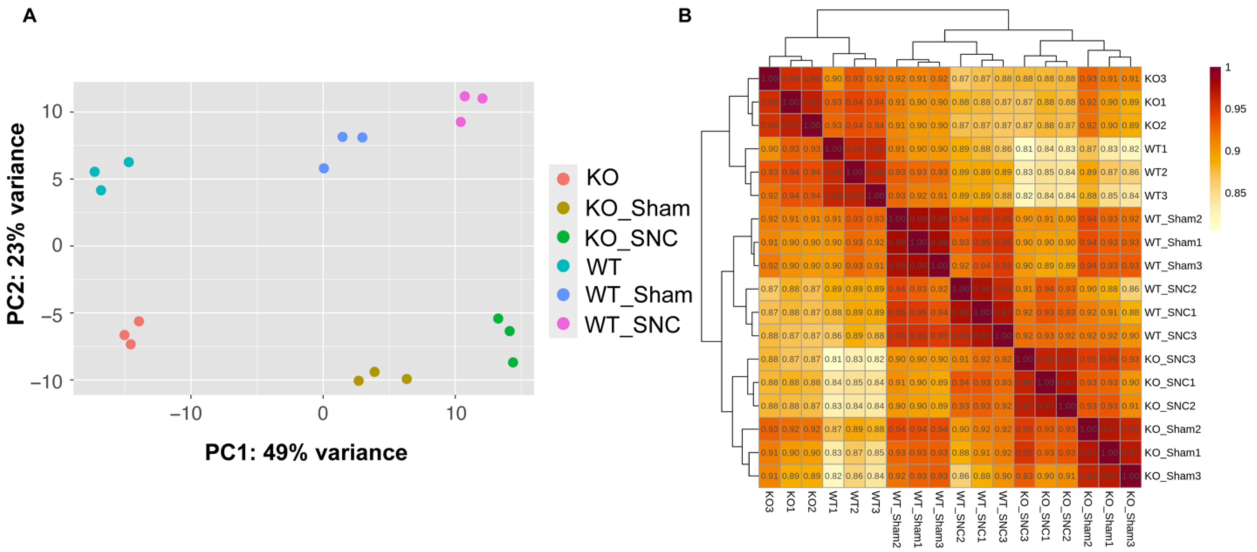


**Supplementary Figure 6. Principal component and correlation analysis.** Principal component analysis (PCA) **(A)** and Spearman correlation analysis **(B)** of transcripts per kilobase million (TPM) for promoter regions (TSS+/-1 Kb) of all differentially expressed (DE) genes (abs(FC)>1.5 and FDR<0.05) obtained from six different condition comparisons (KO-vs-WT, WT_SNC-vs-WT_Sham, KO_SNC-vs-WT_SNC, KO_SNC-vs-WT_Sham, KO_SNC-vs-KO_Sham, and KO_Sham-vs-WT_Sham). Each indicated condition has three independent samples.

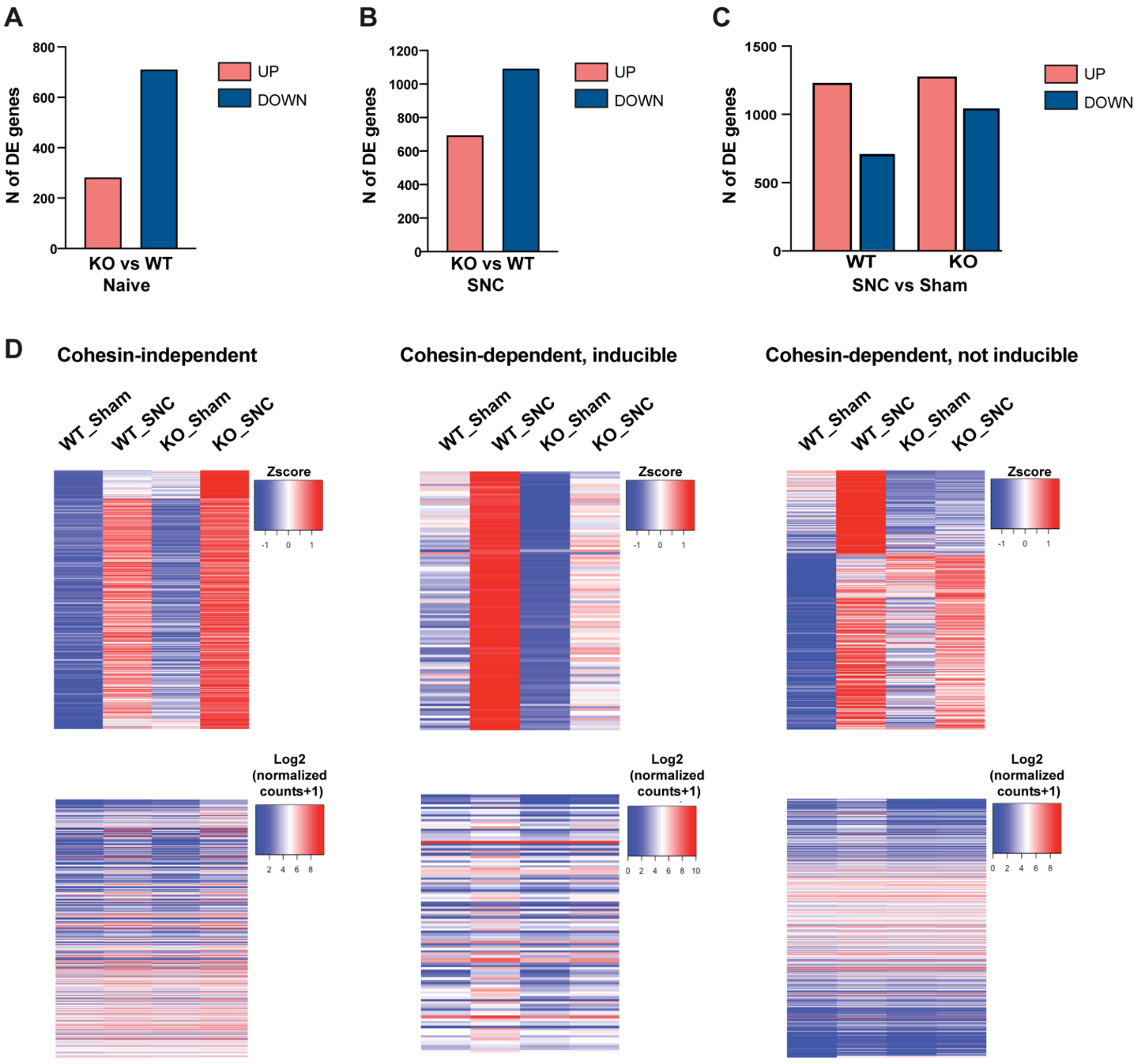


**Supplementary Figure 7. Differentially expressed genes after injury in wt and cohesin-depleted neurons. (A-C)** Bar charts of the differentially expressed (DE) genes in the indicated conditions (*n*=3 independent samples; FDR<0.05). Pink and blue represent upregulated and downregulated genes, respectively. **(D)** Heatmaps of the Z score and expression level of the cohesin-independent and dependent (inducible and non-inducible) genes in the indicated conditions.

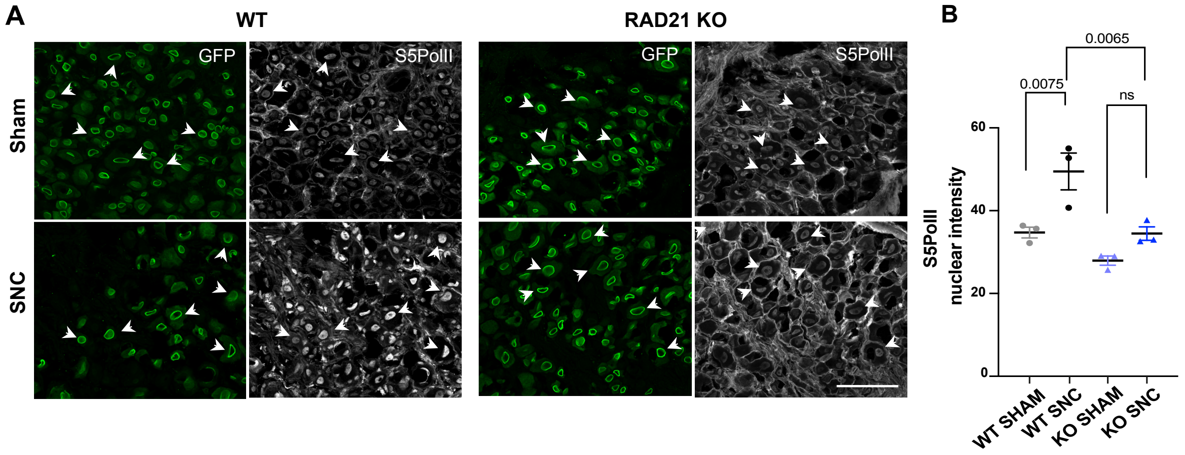


**Supplementary Figure 8. Lack of cohesin impairs injury-dependent PolII activation. (A)** Micrographs showing nuclear GFP (green) and Serine 5 phospho RNA Polymerase II (S5Pol II, grey) expression in DRG neurons 16 days after 4 consecutive daily tamoxifen injections in INTACT-AdvCre (WT) and INTACT-AdvCre-*Scc1*flox/flox (*RAD2*1 KO) mice, 3 days post sciatic nerve crush (SNC) vs Sham. Arrowheads mark nuclear GFP positive neurons. Scale bar, 100 µm. **(B)** Scatter plot of the quantification of S5Pol II nuclear signal intensity (mean ± s.e.m of *n*=3 mice; one-way Anova, Tukey's multiple comparison test).

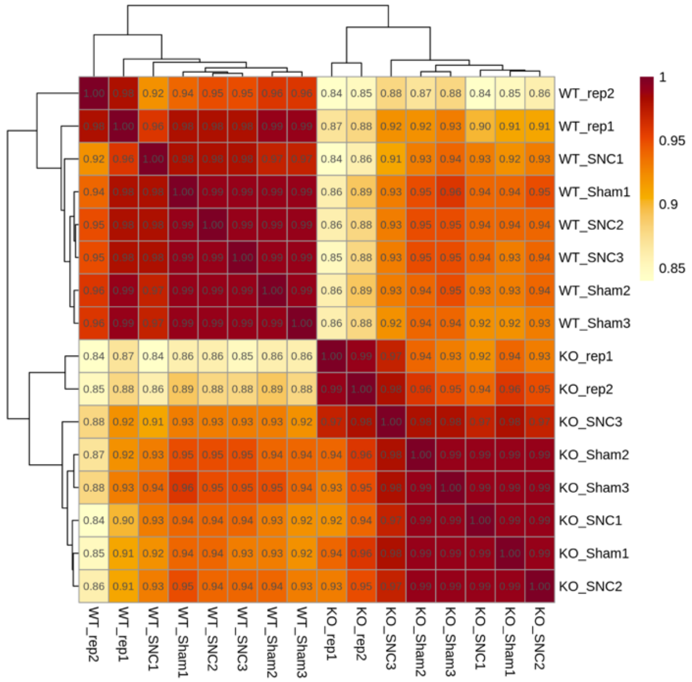


**Supplementary Figure 9. Hi-C reproducibility.** Hi-C reproducibility results from hierarchical clustering of stratum-adjusted correlation coefficients (SCC) of Hi-C contact matrices at 100-Kb resolution in the indicated conditions (*n*=3 or 2 independent samples).

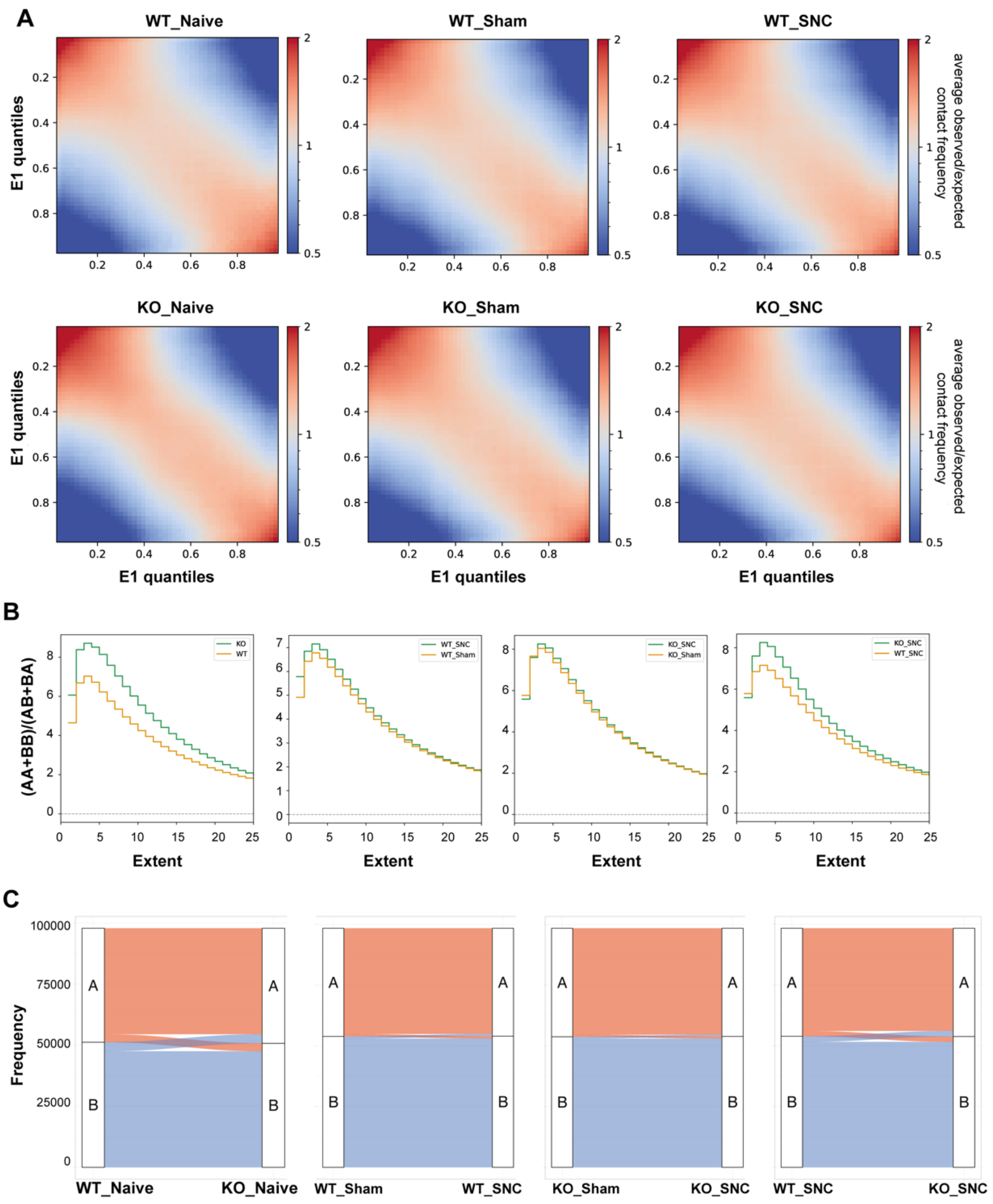


**Supplementary Figure 10. (A) Lack of cohesin does not affect compartimentalization.** Saddle plots of the average observed/expected contact frequency between A and A (bottom-right), B and B (up-left), A and B (bottom-left), and B and A (up-right) compartments in the indicated conditions. **(B)** Saddle strength plots of the ratio of (AA+BB) / (AB+BA) interaction frequency in the indicated conditions. **(C)** Alluvial plots on the whole genome indicating the number of A or B compartment regions that are changed to B or A, respectively, in the indicated conditions.
